# Highly mutated memory cells drive an inefficient secondary antibody response to a variant protein

**DOI:** 10.1101/438184

**Authors:** Richard K Tennant, Barbara Holzer, John Love, Elma Tchilian, Harry N White

## Abstract

A powerful vaccine against mutable viruses might induce memory antibodies that either strongly bound antigenic variants or that could rapidly undergo secondary affinity maturation to achieve this. We have recently shown after secondary immunization of mice with a widely variant protein (Burton et al. 2018) that IgM+ memory B-cells with few mutations supported an efficient secondary germinal centre (GC) and serum response, superior to a primary response to the same protein. Here, boosting with more closely related proteins produced a GC response dominated by highly mutated B-cells that failed, not efficiently improving serum avidity even in the presence of extra adjuvant, and that was worse than a primary response. This supports a hypothesis that over certain antigenic differences, a cross reactive, mutated, memory B-cell compartment can be an impediment to affinity maturation.

## Introduction

Globally important diseases such as Dengue and Influenza spread across populations as epidemics. While strain-specific immunity can be induced by vaccination or infection, the continuing appearance of variant viruses presents a major challenge to vaccine design.

Foreign antigens stimulate the formation of memory B-cells that have undergone somatic hyper-mutation (SHM) of their antibody genes followed by selection in germinal centres (GCs) for antibody affinity maturation(MacLennan et al. 1997). In this manner protective immunity is established against subsequent exposure to identical pathogens. What is poorly understood, but critical for the understanding of immune responses to mutable pathogens, is how memory B-cells respond to variant antigens.

We have recently shown that antibody memory responses to variant/heterotypic Dengue envelope proteins involve secondary germinal centres (GCs) with a higher proportion of IgM+ B-cells that have fewer VH mutations (Burton et al. 2018) compared to the memory response to the homotypic antigen. These observations provide support for the long speculated idea that ‘lower’ layers of the B-cell memory compartment, with less SHM, could furnish secondary responses against variant antigens, as the antibodies may be more tolerant of antigenic variation (Herzenberg et al., 1980; Pape et al., 2011; Kaji et al., 2012).

A complication of sequential infection by certain combinations of variant viruses, however, is that immune responses to the second virus can be compromised, through a process sometimes termed original antigenic sin (Francis, 1960) or antigenic seniority(Henry et al. 2018). We use the abbreviation AS for both.

In AS, cross-reactive memory-derived antibodies are induced which have a lower avidity, and may increase pathology, as they are non-neutralising or enhance infectivity (Halstead et al. 1977, Halstead et al. 1983, Kim et al. 2009, Midgley et al. 2011). Such effects of AS may facilitate Dengue epidemics(Adams et al. 2006), could enhance Zika pathogenesis in Dengue endemic areas (Bardina et al. 2017) and may be induced by vaccination(Screaton et al. 2015).

While it is not surprising that a similar antigen induces a cross-reactive memory response rather than a naïve response, due to increased numbers of memory cells and their lower activation thresholds(Good et al. 2007, Good et al. 2009), a critical question in AS is why such memory responses then fail to rapidly evolve higher affinity to variant antigens through a secondary GC reaction, mitigating pathogenesis.

There is a theoretical model that addresses this question(Deem et al. 2003). It predicts that over certain antigenic differences, antibody memory responses to a variant antigen are worse than a primary response; *i.e*., that memory responses can be compromised in their ability to evolve toward certain antigens. Using sequential immunization with variant Influenza haemagglutinins and analysis of serum and GC responses, we have tested predictions of this hypothesis and provide evidence that it is correct. We show that an inefficient secondary response to a variant haemagglutinin is associated with recruitment of highly mutated memory B-cells to the GC, altered patterns of SHM, collapse of the GC reaction and poor serum avidity improvements.

Taken together with our recent report of memory responses to heterotypic Dengue E-proteins, these results provide important measures of success and failure in anti-variant memory B-cell responses that should be of value in the design of vaccines that elicit the broadest protection against viral variation.

## Results

### Primary and secondary response to PR8 HA

Firstly we sought to define the serum and GC response after homotypic antigen prime and boost. Mice were primed with PR8/34 (PR8) HA1 with adjuvant and boosted 38 days later with soluble PR8 HA (see Methods for explanation of rationale).

PR8 HA1 priming induced an anti-PR8 HA IgG titre by day 17, increasing to day 44 (Figure 1A). Boosting increased the titre modestly (by two-fold) but significantly by day 17 post boost.

Time points were chosen to assess recent recruits to GCs (day 6) and then after selection had been established (day 17)(Victora 2014). Levels of GC B-cells rose from a background of 0.18% to 0.75% by day 6 after priming, remained elevated to day 17, and fell to background by day 44 (Figure 1 B, C). Boosting induced GC B-cells to a higher level: 1.5%, by day 6, which reduced but not significantly by day 17 (Figure 1C).

VH mutations increased by day 17 after priming, consistent with active SHM and affinity maturation (Figure 1D). Boosting induced GC B-cells with a higher level of VH mutation at day 6 (median=3) than seen at day 6 after priming (median=1.5), consistent with a response mostly from memory B-cells focused on the HA1 head region. GC B-cells then accumulated further mutations by day 17 after boost (median=6), consistent with further affinity maturation.

**Figure 1.**
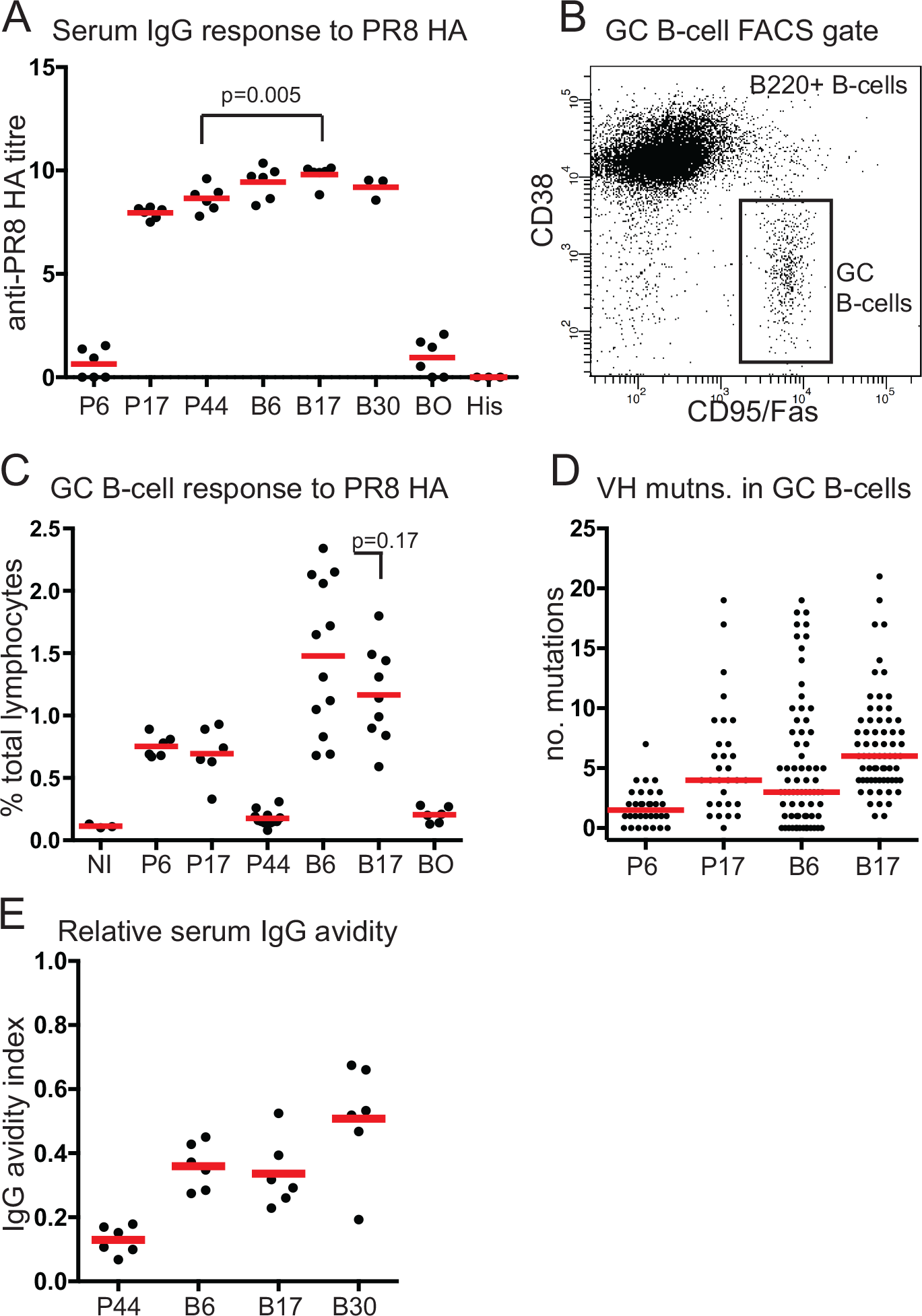
Primary and secondary response to homotypic PR8 HA. **A**, Anti-PR8 HA serum IgG response. Bar shows mean. Px, x days since priming; Bx, x days since boosting; BO, adjuvant-only primed, PR8 HA boosted, analysed day 6; His, B6 serum sample reactivity with irrelevant C-terminal His-tagged protein (*B. Clausii* pdxR). **B**, FACs gating strategy. **C**, GC B-cell response to PR8 HA. Bar shows mean. NI, not immunised; other labels as panel A. **D**, VH mutations in single sorted GC B-cells. From n=3 mice (Px) and n=6 mice (Bx). Bar shows median value. Labels as panel A. **E**, Relative serum IgG avidity for PR8 HA. Bar shows mean. Labels as panel A.

Analysis of the relative avidity of serum IgG (Figure 1E) showed a rapid increase by day 6 after PR8 HA boosting, consistent with preferential stimulation of the highest affinity memory cells by the homotypic boost, and the subsequent immediate differentiation of many of them into antibody-secreting cells. There was no detectable avidity increase by day 17 but a further increase was detected by day 30.

### A failed secondary response to heterotypic HA

After PR8 HA1 priming, we then performed boost immunisations with Bris59 HA which has 82% identity with PR8 over the HA1 region.

Figure 2A shows that despite a robust response by day 6, up to just over 1.0%, the GC response had fallen back by day 17 to 0.3%, close to the pre-boost background level of 0.18%. This contrasted with the sustained response seen after PR8 HA boosting, and the responses seen 17 days after boosting with heterotypic Dengue proteins, which were still four to eight-fold above background at day 17 (Burton et al. 2018).

Analysis of mutations in all Bris59 HA boosted GC B-cells showed an increase, from a median of 3 at day 6, to 6 at day 17, the same profile as the PR8 HA boost (Figure 2B). This figure also shows that levels of mutation at day 6 in IgM+ GC B-cells were the same (median=3) in PR8 and Bris59 HA boosted groups. With the previously reported response to heterotypic Dengue proteins, early GC B-cells had fewer mutations, compared to the homotypic boost (Burton et al. 2018).

**Figure 2.**
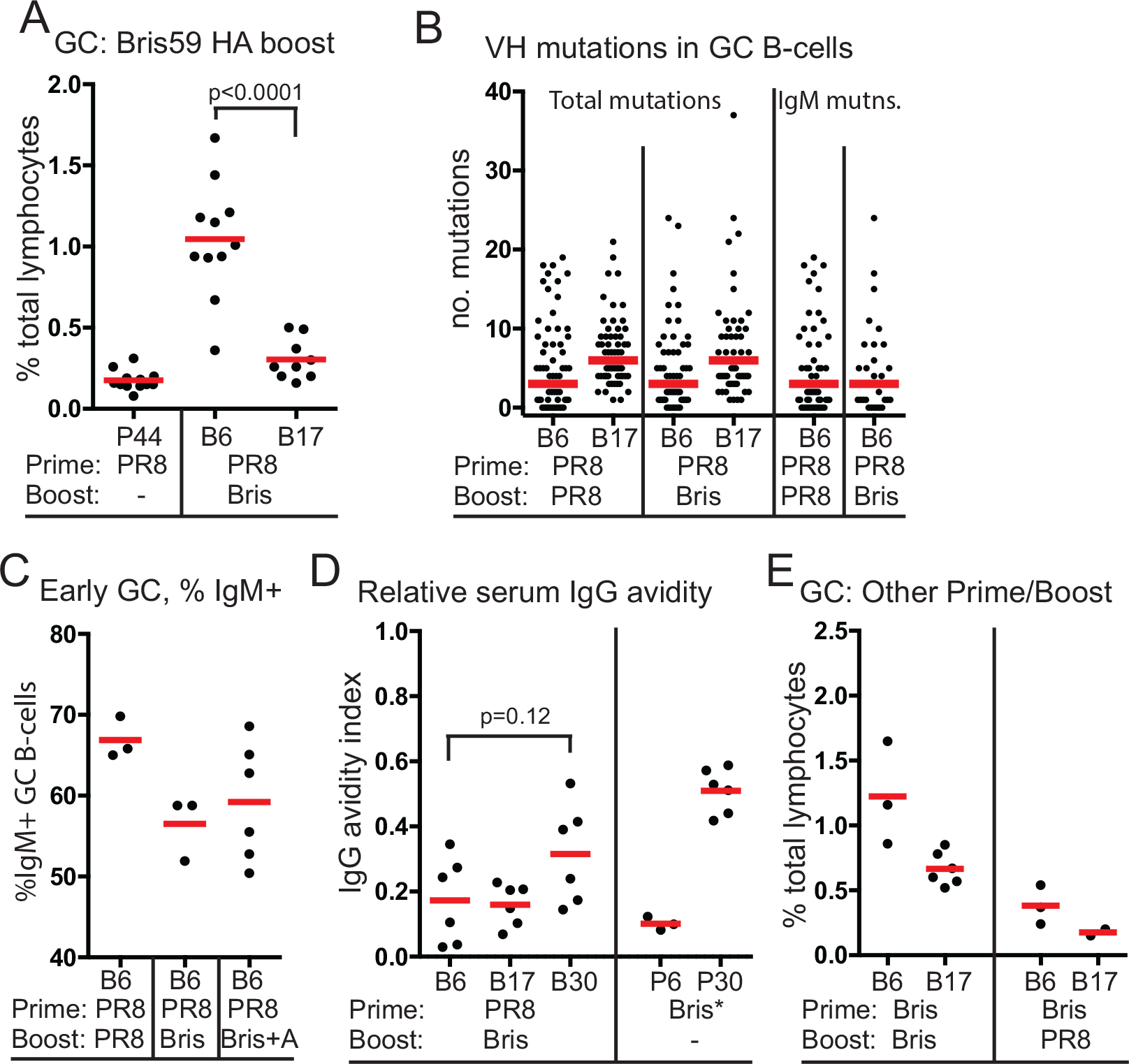
Secondary responses to variant HA boosting. **A**, GC B-cell response to Bris59 HA boost. Bars show mean. First two data sets reproduced from panel A for comparison. X-axis labels: Px, x days since priming with antigen indicated below; Bx, x days since boosting with antigen indicated below. Bris, Bris59 HA. **B**, VH mutations in single sorted GC B-cells. From n=6 mice in each group. Bar shows median. Bx, days since boost with antigen indicated below. First 2 data sets reproduced from Figure 1D for comparison. **C**, Proportion of IgM+ GC B-cells. X-axis labels as for panel B. Bris+A, Bris59 HA including adjuvant. **D**, Relative serum IgG avidity for Bris59 HA. Bars show mean. X-axis labels as for panel B. Note: The Bris* prime was performed with Bris59 HA, not HA1, to allow appropriate comparison with Bris59 HA boosted groups. **E**, GC B-cell levels after homotypic Bris59 HA1/Bris59 HA boost and heterotypic Bris59 HA1/PR8 HA boost. Labels as for panel B.

Further, here, at day 6 after Bris59 HA boosting, there was a lower proportion of IgM+ B-cells (Figure 2C) compared to the homotypic PR8 boost, again contrasting with the Dengue protein responses where heterotypic antigen boosting induced GCs with a higher proportion of IgM+ cells. This indicates a GC response of a different quality, involving greater proportions of more mutated IgG expressing cells, as compared to previously observed heterotypic GC responses. There was no significant increase in the relative avidity of serum IgG detected by day 30 after Bris59 HA boost, to 32% (Figure 2D). This rise was less than seen after a Bris59 HA prime, up to 51% at day 30 (Figure 2D), and heterotypic Dengue-4 envelope-protein boosting, up to 53% by day 32 (Burton et al. 2018). To test whether Bris59 HA is not just a weaker antigen, we performed a homotypic Bris59 HA1/Bris59 HA prime-boost. We also did the reverse heterotypic Bris59 HA1/PR8 HA prime-boost. The homotypic Bris59 prime-boost induced similar levels of early GC B-cells (1.2%; Figure 2E) to the homotypic PR8 prime-boost (1.5%; Figure 1C), which then declined slightly by day 17 to 0.7%, and two-fold higher serum IgG titres (Figure 3A) compared to the homotypic PR8 prime-boost (Figure 1A), suggesting its antigenicity is similar and not explaining the drop in GC B-cell levels after the heterotypic PR8 HA1/Bris59 HA prime-boost. Interestingly, the reverse heterotypic Bris59 HA1/PR8 HA boost did not induce high levels of early GC B-cells (Figure 2F), showing that this type of heterotypic boost response is antigen order dependent.

### Effect of adjuvant on the heterotypic Bris59 HA boost response

Adjuvants containing TLR ligands can increase GC responses(Kasturi et al. 2011) and squalene oil-in-water adjuvants can mitigate AS(Kim et al. 2012). We hypothesized that a squalene oil-in-water adjuvant may improve the function of the GCs undergoing the boost response to Bris59 HA. This adjuvant included a TLR-4 ligand, MPL, which due to the ability of MPL/LPS to stimulate B-cells(Anderson et al. 1972), may improve recruitment of less mutated memory cells which we have shown are associated with successful heterotypic secondary responses (Burton et al. 2018).

Adjuvanted Bris59 HA boosts induced a nearly four-fold increase in IgG titre at day 17, above that of the non-adjuvant boost (Figure 3A), and induced sustained GC B-cell levels, rising to 1.2% of lymphocytes by day 6 and only falling to 0.9% by day 17 (Figure 3B). At day 30, however, the IgG titre was only slightly more in adjuvanted Bris59 HA boosted mice compared to Bris59 HA primed mice (Figure 3A). Considering pre-existing IgG titre prior toBris59 HA boosting, the increase in titre may be less.

**Figure 3.**
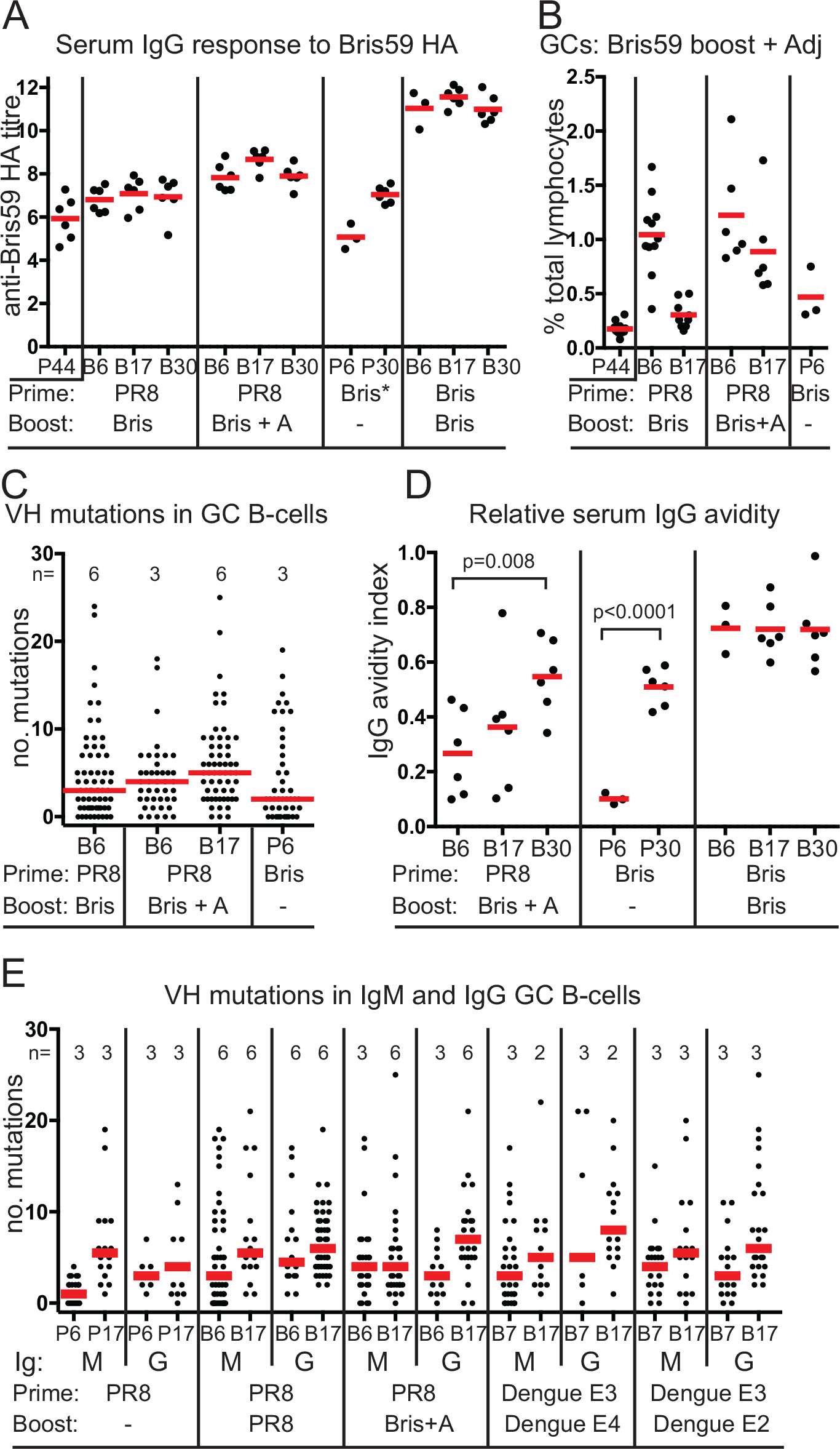
Effect of adjuvant on heterotypic Bris59 HA boost. **A**, Anti Bris59 HA serum IgG titres after Bris59 HA boosting. Bars indicate mean. X-axis labels: P44, PR8 HA1 prime only, day 44. All other labels: Px, x days since priming with antigen indicated below; Bx, x days since boosting with antigen indicated below. Bris + A, Bris59 HA including adjuvant. Note: The Bris* prime was performed with Bris59 HA, not HA1, to allow appropriate comparison with Bris59 HA boosted groups. **B**, GC B-cell levels after Bris59 HA boost with and without adjuvant. First three groups reproduced from Figure 2 panel B. Bars indicate mean. X-axis labeling as for panel A. **C**, VH mutations in GC B-cells. Number of mice in each group indicated as n=. Bars indicate median. X-axis labels as for panel A. **D**, Relative serum IgG avidity for Bris59 HA. Bars indicate mean values. Labels as panel A. **E**, VH mutations in IgM+ and IgG+ GC B-cells. Number of mice in each group indicated as n=. Bars indicate median. X-axis labels as for panel A.

GC B-cells induced six days after the adjuvanted boost had similar if not greater levels of VH mutation (median=4) compared to the non-adjuvanted boost (median=3; Figure 3C), and an equivalent proportion of IgM GC B-cells (Figure 2C), implying that no extra naïve, or naïve-like memory cells, were recruited to GCs by adjuvant. By day 17 median VH mutations in this group had only increased by one (Figure 3C). GC B-cells 6 days after Bris59 HA priming had lower levels of VH mutation (median=2), consistent with a primary response.

The adjuvanted Bris59 HA boost induced a significant increase in serum avidity from day 6 to day 30, but only to levels comparable to those seen with the Bris59 HA prime group, being 54% *vs.* 51%, and not approaching the avidity of the homotypic Bris59 HA1/Bris59 HA prime–boost which was 72% (Figure 3D). Considering the net level of increase in avidity between day 6 and day 30, the Bris59 HA prime appeared more efficient (41% increase) than the adjuvanted heterotypic boost (28% increase).

### Altered selection of VH mutations in Bris59 HA boosted mice

Despite a sustained GC reaction, the adjuvanted Bris59 HA response only supported a slight increase in overall VH mutation (median=+1) between days 6 and 17 (Figure 3C), compared to the PR8 HA1 primary (+2.5), PR8 HA boost (+3) and Dengue-4 envelope protein boost (+4) (Burton et al. 2018). As well as a smaller overall increase in SHM, analysis of the levels of VH mutations separately in IgM and IgG, showed further evidence of altered selection. The primary and secondary responses to PR8 HA and the heterotypic Dengue protein E4 and E2 responses all showed increases in IgM VH mutation of between +1.5 and +4.5 as the GC response matured (Figure 3E). In the adjuvanted Bris59 HA boost response there was no increase in IgM mutations and the largest increase in IgG mutations between day 6 and day 17. The non-adjuvanted Bris59 HA boost response was not included in this analysis as the failure of the GC reaction implies few cells with any level of mutation were successfully selected to day 17.

## Discussion

We have shown how B-cell memory and GC responses to boosting with heterotypic antigens can be inefficient. With the non-adjuvanted heterotypic Bris59 HA boosts, GC B-cell numbers collapsed by day 17 despite ongoing SHM, and serum avidity failed to increase significantly, contrasting with heterotypic Dengue protein boosts in mice. With adjuvant, Bris59 HA boosting induced comparable IgG titres but a smaller increase in serum avidity than the primary response to Bris59 HA. Considering the adjuvanted Bris59 HA boost response initiated with 3.6-fold higher numbers of GC B-cells above background (Figure 3B), it is less efficient than the primary response. Despite providing a larger and sustained GC response, therefore, the presence of a cross-reactive memory B-cell compartment was an impediment to the Bris 59 HA response.

We previously reported that successful heterotypic secondary responses had early GCs containing higher proportions of IgM+ B-cells with fewer mutations compared to the homotypic boost (Burton et al. 2018). The heterotypic responses to Bris59 HA reported here had early GCs with a lower proportion of IgM+ B-cells than the homotypic boost (Figure 2C), with equivalent or even higher levels of mutation compared to the homotypic antigen boost (Figures 2B, 3C). In the adjuvanted Bris59 HA boost the higher levels of mutation in IgM at day 6 then failed to increase further during the mutation and selection process to day 17 (Figure 3E). We propose that these heterotypic secondary responses to Bris59 HA were compromised because antibody evolvability was reduced. This could be due either to an intrinsic loss of evolvability in mutated V genes, other constraints in the memory antibody repertoire, or for GC dynamical reasons such as increased SHM and class switching affecting GC B-cell longevity and fate (Gitlin et al. 2016).

The model of Deem and Lee (Deem et al. 2003) provides both a hypothesis for AS, and for heterotypic memory responses in general. By modeling the evolution of memory-derived antibodies, they predict that over a certain window of antigenic difference, memory responses to variant antigens can be worse than primary responses to the same antigen due to constraints in the antibody fitness landscape. This is despite memory B-cells initiating with a higher affinity than naïve cells for variant antigen, which also explains their dominance of the recall response. If antibody evolvability was lost, memory B-cells would be recruited to GCs, SHM would occur, but the GC reaction and affinity maturation would fail. This is what we observe with the non-adjuvanted Bris59 HA boost.

The Deem model assumes that any memory B-cells stimulated by homotypic and heterotypic boosts are equally mutated. It seems the HA1 region of the Bris 59 HA protein was sufficiently similar to PR8 HA1 (82%) to do this and induce the most mutated, and more often class-switched, anti-PR8 HA1 memory cells, which were then inefficient at further improving affinity for Bris 59 HA. This contrasts, in particular, with the Dengue E3 Prime/E4 boost response, where the antigen pair were only 63% identical and the secondary GC were dominated by IgM+ B-cells with very few mutations that then provided for efficient further affinity maturation (Burton et al. 2018), something not predicted by the Deem model.

The model also assumes antigenic difference is uniform, unlike complex antigens with differently variant epitopes. The inefficient GC response we observe here, therefore, is a plausible *in vivo* outcome for a process involving B-cell clones of varying evolvability. Such differences in antibody evolvability could also explain why particular epitopes become immunodominant in secondary responses (Victora and Wilson 2015).

TLR ligands and adjuvants can increase GC responses (Kasturi et al. 2011) and mitigate AS (Kim et al. 2012). Adjuvants could support survival of less fit B-cell variants allowing rare multiple mutations to improve fitness. In the adjuvanted Bris59 HA boost response, that shows a persisting GC reaction, there is a low overall increase in SHM of +1 (Figure 3C) compared to +3 in the non-adjuvanted homotypic boost (Figure 1D), with no increase in IgM mutations but the largest increase in IgG mutations, between day 6 and day 17 (Figure 3E). If beneficial mutations are very rare, then most mutations will result in cell death, so extra mutation levels may be lower, as we observe overall, and also in the IgM+ population. Finally, if an increased number of mutations is necessary to circumvent the constraints of the antibody fitness landscape, successfully selected cells would be expected to have a greater increase in mutation, and this is what we observe in the IgG+ pool between day 6 and day 17.

There are also other reasons why mutated memory antibodies may be less evolvable. They could have lost hotspots for the *aicd* mutator, lost codons predisposed to non-conservative substitutions (Hershberg and Shlomchik 2006) and despite being diverse at the sequence level may be more restricted in V-gene diversity so have more constrained evolutionary trajectories.

The inefficient response to Bris59 HA could be due to reduced T-cell help for variant antigens. T-cell help is necessary for memory B-cell responses to viral proteins (Hebeis et al. 2004), so the robust early GC response to Bris59 HA boosting shows that help was not impaired at this point. Further, we have shown with heterotypic Dengue proteins with only 63% sequence homology, that *in vitro* T-cell re-stimulation was as efficient as with the homotypic protein (Burton et al. 2018), demonstrating that antigens with large overall sequence differences can still efficiently stimulate memory T-cells. The PR8/Bris59 HA1 sequence homology is 82% so we do not consider reduced T-cell help was a major factor here. The response to Bris boosting with adjuvant, with robust and sustained GC activity, also supports this conclusion.

That certain antibody memory responses can have reduced efficiency has important implications for understanding viral disease pathogenesis. It is plausible that epidemic strains of virus could appear with the appropriate degrees of antigenic difference to exploit this phenomenon, as has been previously modeled (Adams et al. 2006). Further, recently proposed strategies to generate broadly neutralizing antibodies propose sequential immunisation with antigens of relatively large mutational distances (Shaffer et al. 2016) and it is important to determine how these might induce compromised memory responses.

## Materials and Methods

### Mice, immunisation, antigens

All HA and HA1 proteins were from Sino Biologicals; expressed in 293 cells. Female 8–11-week BALB/c mice were from Charles River, U.K. Most primary immunisations: i.p.: 25 μg PR8/34 haemagglutinin (HA)1 alum precipitated with 2 × 10^7^ heat-killed *B.pertussis*. Secondary immunisations: i.p., 25 μg HA protein in PBS, or with Sigma Adjuvant System (Sigma). Blood samples collected under terminal anaesthesia, and spleen cells after subsequent sacrifice. Rationale for HA1 priming/HA boost was to focus the response away from the conserved stem and on to the variable epitopes of the head, as we were interested in responses to variant regions. Sequential immunization with variant whole HAs focuses the response onto the conserved stem region (Ellebedy et al. 2014). As the C-terminal region of HA1 is also more conserved between strains than the rest of the head region, we wanted to avoid antibodies focused on this areas too, and to change the context of the C-terminal His-tag between prime and boost, to reduce confounding sequence similarities. Bris and PR8 HAs had 0.1 and 0.34 endotoxin U/ug, which at doses used is 2.5% and 9% of a dose known not to cause a physiologic response in mice (Copeland et al. 2005). HA1/HA strains(NCBI accession number https://www.ncbi.nlm.nih.gov/protein): H1N1: PR/8/34(ABD77675.1); Brisbane/59/07(ACA28844.1). All animal experiments were performed under UK Home Office license PPL 30/3089 with permission from the University of Exeter local animal welfare ethical review board.

### ELISA

Standard protocol with: 1 μg/ml coating protein in bicarbonate buffer, PBS/0.05% Tween-20 (Sigma; PBST) washes, PBST/2% BSA (Sigma) block, serum dilutions in PBST/1.0% BSA, alkaline phosphatase conjugated goat anti-mouse IgG (Sigma) second layer, developed with pNPP substrate (Sigma). End-point titre values plotted are log2 of 1/(end point dilution x 100), each unit increase represents a doubling of titre.

### Urea avidity ELISA

From Puschnik *et al.* 2013. Standard ELISA, then after serum step, 1X wash, 10min. incubation with 7M Urea/PBST, 2X wash, then standard protocol. Avidity index: 7M urea readings/readings from PBST-only treatment, after subtraction of background.

### Flow cytometry

RBC-depleted spleen cells were incubated with Fc-block (BD) then allophycocyanin anti-B220, BV421 anti-CD38 and PE anti-CD95/Fas (BD), anti-IgM (eBioscience), following standard protocols.

### B-cell antibody sequencing

Single GC B-cells were sorted into 10 μl chilled 10mM Tris pH 8.0, 1 U/μl RNAsin (Promega) and frozen at -80°C. One-Step RT-PCR (Qiagen) was done with Tiller et al. 2009 primer sets, for 50 cycles annealing at 53.6°C. Heavy-chain second-round PCRs with 2 μl first-round PCR, Tiller et al. 2009 primers, with Taq (Qiagen) for 50 cycles annealing at 56°C. Products were sequenced using primer 5’MsVHE(Tiller et al. 2009) which leaves part of 5’ of FR1 unsequenced, hence FR1 was not analysed. VH sequences were analysed using IMGT V-Quest platform. Sequences: Supplemental Table 1.

### Statistics

Three mice were randomly allocated to cages. For greater sample sizes treatments were independently replicated. Where t-test was applied, data points were analysed for equality of variance, and where violated were subject to a two-tailed test for unequal variance, otherwise a two-tailed test for equal variance.

## Acknowledgements

We are grateful to the late Michael Neuberger for early critical discussions. Thanks to Patrick Wilson and Christian Busse for advice on single cell antibody PCR and James Cresswell for advice on statistics.

